# Integrative enhancer discovery identifies functional enhancer dependencies in pediatric acute myeloid leukemia

**DOI:** 10.64898/2026.08.18.745242

**Authors:** Leah Schüler, Robert Winkler, José Gonçalves-Dias, Konstantin Schuschel, Hasan Issa, Lonneke Verboon, Xiaoyan Wei, Ronay Cetin, Yves Matthess, Manuel Kaulich, Stefan Hüttelmaier, Raj Bhayadia, Dirk Heckl, Jan-Henning Klusmann

**Author notes:** Contact information for correspondence: Prof. Dr. Jan-Henning Klusmann. These authors contributed equally to this work (Dirk Heckl, Jan-Henning Klusmann).

## Abstract

Pediatric acute myeloid leukemia (AML) is driven by aberrant transcriptional programs sustained by poorly defined cis-regulatory mechanisms. To systematically identify functional enhancer dependencies, we developed an integrative enhancer discovery strategy that combines H3K27ac CUT&Tag profiling, enhancer-associated transcription, and CRISPR interference (CRISPRi) screening. By leveraging enhancer-associated transcription to prioritize candidate regulatory elements, we identified 321 leukemia-associated enhancers for functional interrogation. This approach uncovered the hematopoietic *MYB* enhancer (H-ME) within the HBS1L-MYB-AHI1 locus as a critical regulator of leukemic growth. H-ME repression reduced chromatin accessibility and active histone marks at the *MYB* promoter, suppressed *MYB* expression, and induced differentiation-associated transcriptional programs. In contrast, selective depletion of the enhancer-associated transcript had no effect on *MYB* expression or leukemic proliferation, demonstrating that enhancer activity resides within the underlying regulatory DNA element rather than its mature RNA product. H-ME exhibited preferential activity in megakaryocytic leukemia, and its perturbation impaired leukemic growth in primary patient-derived models in vitro and in vivo. Together, our findings establish an integrative framework for the systematic discovery of functional enhancer dependencies and identified H-ME as an RNA-independent regulator of MYB in pediatric AML.

## Introduction

Acute myeloid leukemia (AML) in children remains an aggressive malignancy with relapse as the major cause of treatment failure.^1–4^ While therapeutic advances have improved outcomes, further progress will depend on identifying oncogenic dependencies beyond recurrent coding mutations since genetic alterations do not fully explain how leukemic transcriptional programs are established and maintained. Increasing evidence indicates that aberrant gene-regulatory circuits and epigenetic dysregulation are central determinants of leukemic cell identity, differentiation arrest, and disease persistence.^5–11^ Defining the cis-regulatory elements that sustain these transcriptional programs may therefore reveal novel biological dependencies and therapeutic vulnerabilities.

Enhancers are key cis-regulatory elements that orchestrate lineage-specific gene expression by integrating transcription factor binding with chromatin architecture. Active enhancers are typically marked by H3K27ac and communicate with their target promoters through dynamic chromatin interactions.^12^ In leukemia, enhancer landscapes are extensively rewired by oncogenic mutations, altered chromatin states, and aberrant transcription factor activity, resulting in sustained expression of genes that promote malignant self-renewal and block differentiation.^13,14^ While genome-wide epigenomic profiling readily identifies thousands of putative enhancer elements, only a minority are functionally required for leukemia maintenance. Distinguishing these functional enhancer dependencies from permissive chromatin therefore remains a major challenge.

One feature of active enhancers that may facilitate their systematic prioritization is enhancer-associated transcription. Active enhancers frequently produce enhancer RNAs or longer non-coding transcripts, reflecting ongoing transcriptional activity at regulatory elements.^15,16^ Unlike chromatin marks alone, enhancer-associated transcription provides additional evidence that an enhancer region is actively engaged within the transcriptional network.^17^ We therefore hypothesized that integrating enhancer-associated transcription with H3K27ac profiling would enable systematic prioritization for functional screening of leukemia-associated enhancers.

Among key transcriptional regulators in hematopoiesis, MYB plays a central role in maintaining progenitor cell identity and controlling lineage commitment.^18,19^ Dysregulated MYB activity has been implicated in leukemogenesis across multiple AML subtypes, where it supports proliferation and blocks differentiation.^20–22^ The HBS1L-MYB locus contains multiple distal regulatory elements that control *MYB* expression in hematopoietic cells, including enhancers implicated in erythroid differentiation and fetal hemoglobin regulation.^23–25^ More recent studies have further highlighted enhancer-mediated and enhancer-RNA-associated mechanisms of *MYB* regulation.^26–28^ However, whether enhancer-associated transcription can be exploited to systematically identify functional enhancer dependencies in pediatric AML has not been investigated.^29^

Here, we developed an integrative strategy to identify functional enhancer dependencies in pediatric AML by combining H3K27ac CUT&Tag profiling, transcriptomic analysis, and CRISPR interference (CRISPRi) screening. By leveraging enhancer-associated transcription to prioritize candidate regulatory elements, we identified the hematopoietic *MYB* enhancer (H-ME) within the HBS1L–MYB–AHI1 locus as a critical dependency in megakaryocytic leukemia. We demonstrate that H-ME maintains *MYB* expression through an RNA-independent mechanism, define its essential transcription factor circuitry, and establish its functional relevance in primary patient-derived xenograft models. Together, our findings establish an effective framework for systematic enhancer discovery and identify a context-dependent regulatory vulnerability in pediatric AML.

## Materials and Methods

### CRISPR library screens

The single guide RNAs for CRISPRi screenings were designed using the CRISPick tool from the Broad Institute (https://portals.broad institute.org/gppx/crispick/public) based on the Azimuth 2.0 algorithm. Four sgRNAs were chosen for each gene, resulting in a library of 1221 single guide RNAs. For the saturation mutagenesis screen, libraries comprising 1705 sgRNAs targeting the H-ME region and 23619 sgRNAs spanning the HBS1L-MYB-AHI1 locus, respectively, were used. sgRNAs libraries were cloned into the pLenti_mpx_SpCas9 vector (#189632, Addgene, Watertown, MA, USA).^30^ Non-targeting control sgRNAs were designed against the firefly luciferase gene and were cloned into the single guide vectors (Addgene #89395 or #100864).^31,32^ Details on performance of the screenings are provided in the supplementary methods.

### Human participants

Experiments on human material were performed in accordance with the Declaration of Helsinki and have been approved by the Goethe University Ethics Committee. Informed consent was obtained from all human participants or custodians. Human CD34+ HSPCs were isolated from the mobilized peripheral blood of anonymous healthy donors. AML samples were provided by the Berlin-Frankfurt-Münster Study Group (AML-BFM-SG, Germany), the Department of Hematology, Hemostasis, Oncology and Stem Cell Transplantation (Hannover Medical School, Germany) and the Department of Pediatrics (University Medicine Frankfurt, Germany). Please refer to Supplementary Table 1 and 2 for patient characteristics, including sex and age. Information about gender and patient characteristics was not collected for the anonymous healthy donors.

### ATAC-Seq and CUT&Tag

For epigenetic profiling using CUT&Tag and ATAC sequencing, cells were sorted after transduction with the sgRNA virus and freshly processed. ATAC-Seq and CUT&Tag was performed as previously described in Verboon et al.^33^ Used antibodies are stated in Supplementary Table 3.

### Chromatin Immunoprecipitation (ChIP)

ChIP protocol was adapted from Xu et al.^34^ Used antibodies are stated in Supplementary Table 3. qPCR on purified ChIP DNA fragments was performed with primers stated in Supplementary Table 4.

### Mice experiments

Humanized immunodeficient mice^35^ were obtained from Regeneron Pharmaceuticals and bred in the animal facility at the Georg-Speyer-Haus (Paul-Ehrlich-Straße 42-44, Frankfurt am Main, Germany). Immune-deficient mice were irradiated 24 hours before transplantation with 2.5 Gy. After irradiation, the animals received Ciprofloxacin water (0.05 mg/ml) for 24 days to prevent other infections. Transplantation of 1 × 10^6^ cells was performed intravenously (i.v.) using a mouse tail illuminator (#MTI, BiosebLab, France). For the competition assay, a 1:1 mix of sgNTC-dTomato and sgNTC-APC or sgHME-APC was transplanted, and mice were sacrificed after the first onset of disease. Bone marrow was isolated, stained with an anti-human CD45-APC-Cy7 antibody (Cat#368516, Biolegend, San Diego, CA, USA) and analyzed by flow cytometry.

### Statistical Analysis and Definitions

Statistical analyses of gene expression data (RNA-seq) were performed using DESeq2. CRISPR-Cas9 screening data were analyzed using MAGeCK to call essential genes, with the exception of the tiling screens, which were analyzed in R using DESeq2. Differences with p < 0.05 were considered significant. Detailed statistics are available in the supplementary methods.

## Results

### Systematic identification of functional enhancer dependencies in pediatric AML

To systematically identify functional enhancer dependencies in pediatric AML, we first established a comprehensive atlas of active regulatory elements using H3K27ac CUT&Tag profiling across normal hematopoietic populations, primary pediatric AML samples, AML cell lines, and patient-derived xenograft (PDX) models. The resulting dataset comprised 83 pediatric AML patient samples, 13 healthy donor samples, and 14 cell lines or PDX models. Consistent with the context-dependent nature of enhancer regulation, unsupervised clustering revealed distinct enhancer landscapes that clearly separated normal hematopoietic populations from molecular AML subtypes (Figure 1A and Supplementary Figure 1A-B).

**Figure 1:**
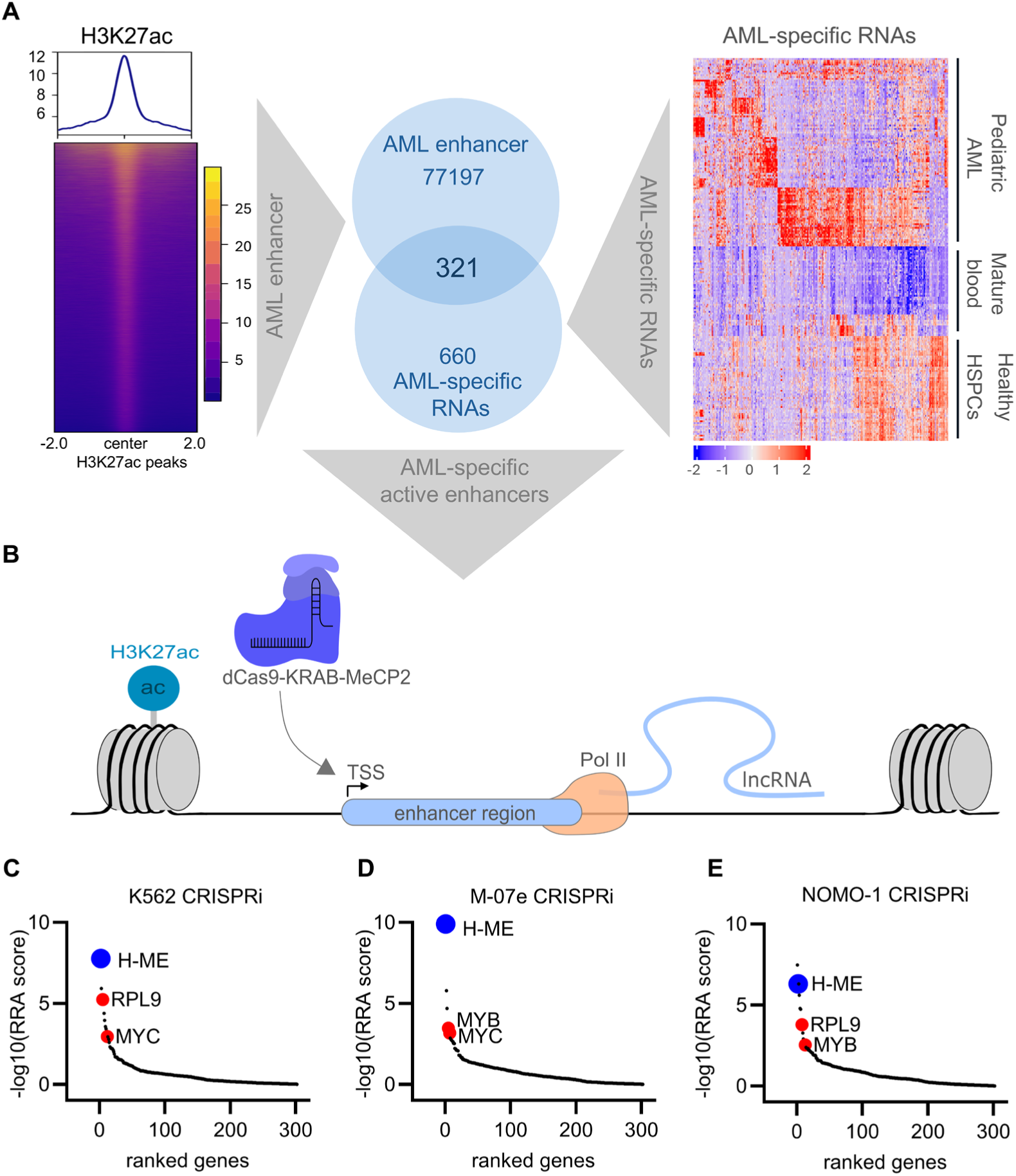
**A** Left Panel: Heatmap of CUT&Tag signal for H3K27ac marks in our hematopoietic database. The database includes 83 patient samples, 13 healthy donor samples and 14 cell lines and PDX samples. Middle Panel: Venn diagram of H3K27ac AML enhancer marks and AML specific RNAs. A total of 321 target genes were selected for library screening. Right Panel: Heatmap of AML specific RNAs. This dataset consisted of 96 healthy donor samples, 106 patient samples and 22 PDX and cell lines. **B** Schematic figure of CRISPRi screening for identification of novel oncogenic DNA-regulatory elements of selected 321 target genes. **C-E** Gene essentiality scores (RRA rank) from MAGeCK analysis of the CRISPRi screens in K562, M-07e and NOMO-1 AML cell line (n = 2 biological replicates each). The enhancer target H-ME is highlighted in blue, and the positive controls RPL9, MYC, and MYB are highlighted in red.

Because chromatin marks alone do not distinguish functional enhancers from permissive regulatory elements, we next integrated enhancer profiling with enhancer-associated transcription.^36^ We therefore analyzed transcriptomic data from 96 healthy donor samples, 106 pediatric AML samples, and 22 AML cell lines and PDX models. Similar to the epigenetic data, enhancer-associated non-coding transcription clearly discriminated normal hematopoiesis from leukemia and identified subtype-specific transcriptional programs (Figure 1A and Supplementary Figure 1C). We hypothesized that regulatory elements supported by both active enhancer chromatin and enhancer-associated transcription would prioritize enhancer candidates for functional screening approaches. We therefore intersected H3K27ac-marked enhancer regions with AML-enriched non-coding transcripts, yielding 321 candidate leukemia-associated enhancers for functional interrogation (Figure 1A).

To directly identify enhancers required for leukemic fitness, we generated a focused CRISPR interference (CRISPRi) library targeting all 321 candidate regulatory elements and performed pooled dropout screens across seven AML cell lines (Figure 1B). Analysis of sgRNA depletion identified several candidate enhancer dependencies, including a regulatory element within the HBS1L-MYB-AHI1 locus (Figure 1C-E and Supplementary Figure 1D-G). Individual validation confirmed that CRISPRi-mediated repression of this region consistently impaired proliferation in K562, M-07e, and NOMO-1 cells (Supplementary Figure 1H).

The identified regulatory element is located within an intron of *AHI1*, immediately adjacent to the *MYB* oncogene. Analysis of publicly available chromatin datasets revealed canonical enhancer features, including strong H3K27ac and H3K4me1 enrichment (Figure 2A). Given its robust phenotype in the CRISPRi screen, its characteristic enhancer architecture, and its proximity to *MYB*, we selected this regulatory element for mechanistic investigation and hereafter refer to it using the recently established nomenclature as the hematopoietic *MYB* enhancer (H-ME).^37^

**Figure 2:**
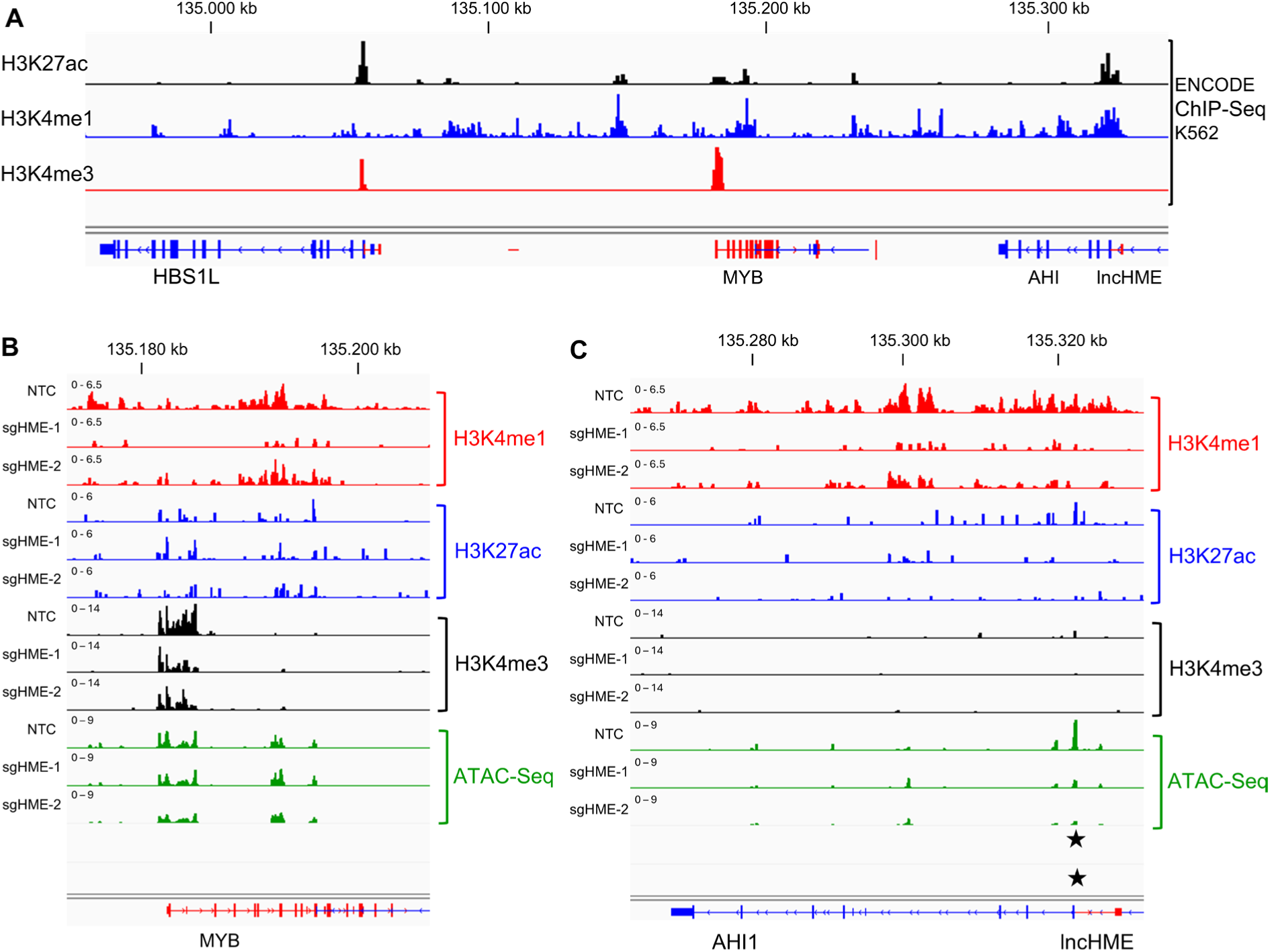
**A** Tracks from the publicly available ENCODE ChIP-Seq data in K562 showing the HBSL1-MYB-AHI1 locus. Histone marks from top to bottom: H3K27ac (black), H3K4me1 (blue) and H3K4me3 (red) are shown in different colors. **B** and **C** Tracks from the ATAC seq and CUT&Tag experiment upon CRISPRi targeting the H-ME enhancer region. CRISPRi sgRNAs are highlighted as stars.

Together, these findings establish an integrative framework for systematically prioritizing functional enhancer dependencies in pediatric AML and identify H-ME as a candidate context-dependent regulator of *MYB* selected for detailed mechanistic investigation.

### H-ME regulates MYB through an RNA-independent chromatin mechanism

To define the mechanism underlying the anti-leukemic phenotype following H-ME perturbation, we first examined expression of genes within the HBS1L-MYB-AHI1 locus after CRISPRi-mediated enhancer repression. While expression of both the enhancer-associated transcript (ENSG00000287094, hereafter referred to as lncHME) and *MYB* was markedly reduced, expression of neighboring genes, including *HBS1L* and *AHI1*, remained largely unchanged (Supplementary Figure 2A-D). These findings indicate that H-ME selectively regulates *MYB* rather than broadly affecting transcription across the surrounding genomic locus.

Because enhancer function is intrinsically linked to local chromatin architecture, we next asked whether H-ME is required to maintain an active chromatin state. ATAC-seq demonstrated reduced chromatin accessibility at both H-ME and the *MYB* promoter following CRISPRi-mediated repression (Figure 2A-C). Consistent with these findings, CUT&Tag revealed a marked reduction of the active chromatin marks H3K27ac, H3K4me1, and H3K4me3, accompanied by accumulation of the repressive mark H3K9me3 at the enhancer (Figure 2B-C and Supplementary Figure 2E-H). Together, these data establish H-ME as a chromatin regulatory element that maintains an active epigenetic state required for *MYB* transcription.

To determine whether reduced *MYB* expression accounts for the observed growth defect, we performed rescue experiments by ectopically expressing *MYB* in CRISPRi-targeted cells (Figure 3A). Restoration of *MYB* substantially rescued the proliferation defect induced by H-ME repression, demonstrating that *MYB* is the functional target of this enhancer (Figure 3B).

**Figure 3:**
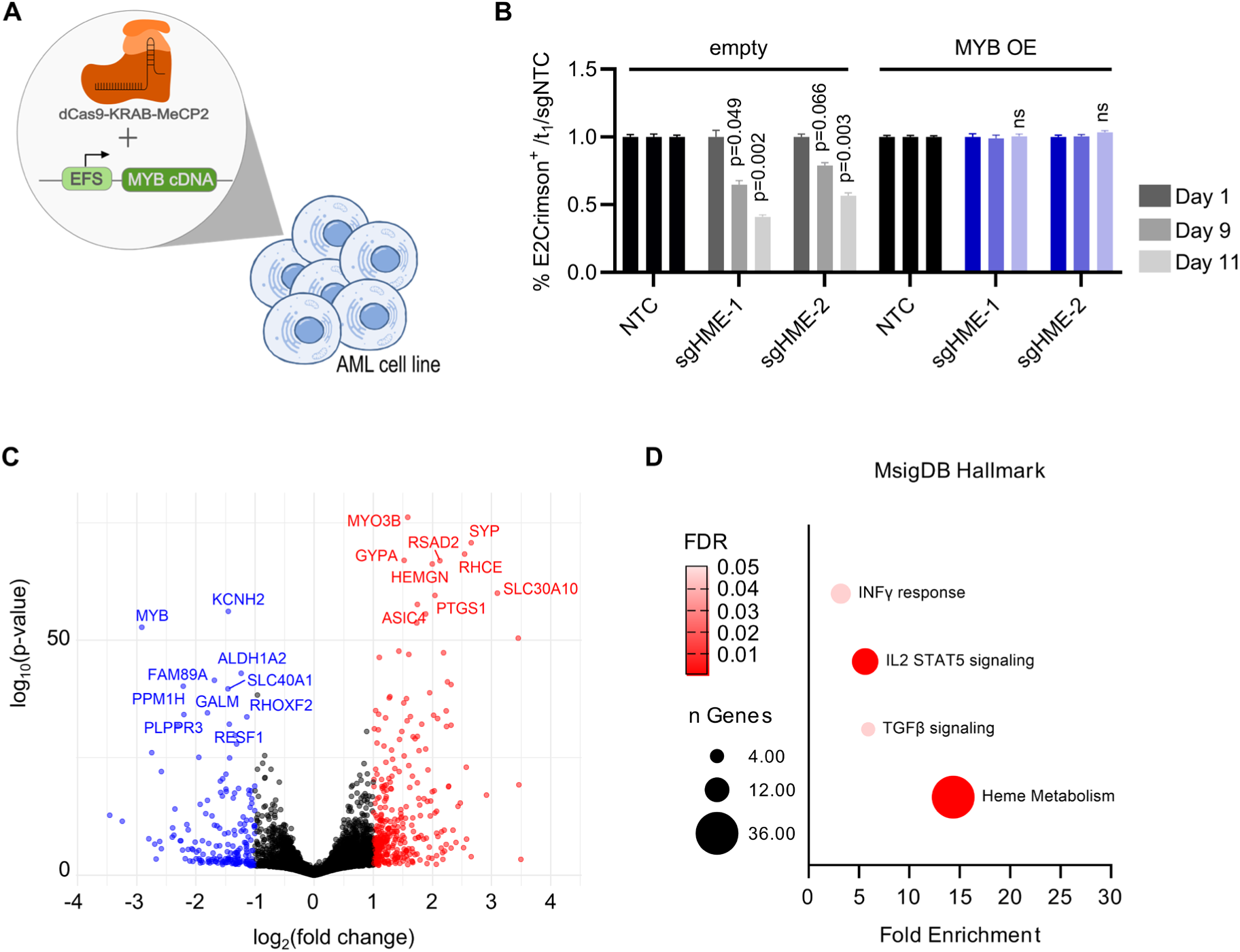
**A** Schematic representation of the rescue experiment. In K562 cells, H-ME was targeted using CRISPRi, while overexpression was achieved with either an empty control vector or MYB cDNA. **B** Fraction of transduced cells in culture over 11 days following CRISPR knockdown of HME with two independent sgRNAs. Cells overexpressing control vectors are shown in grey, and those overexpressing MYB cDNA are shown in blue. Values are normalized to day 1 and shown as mean ± SD; n = 2 independent transductions. Statistics: two-way ANOVA with Dunnett’s multiple comparisons test against the non-targeting control (NTC) at each time point. ns, not significant. **C** Volcano plot depicting differential gene expression following CRISPRi enhancer perturbation. Downregulated genes (p < 0.05, LFC < -1) are shown in blue, and upregulated genes (p < 0.05, LFC > 1) are shown in red. **D** Pathway enrichment analysis of significantly upregulated genes using the Hallmark gene set from MSigDB.

Since H-ME gives rise to the non-coding transcript lncHME, we next investigated whether enhancer function depends on the RNA molecule itself or on the underlying regulatory DNA element. To distinguish between these possibilities, we selectively depleted lncHME using the RNA-targeting CasRx system without perturbing the enhancer DNA (Supplementary Figure 3A). In contrast to CRISPRi-mediated enhancer repression, depletion of lncHME neither altered *MYB* expression nor impaired leukemic proliferation (Supplementary Figure 3B-D). These findings demonstrate that the biological activity of H-ME is mediated by the underlying regulatory DNA element rather than by its associated mature transcript. Thus, the lncHME transcript primarily serves as a marker of enhancer activity rather than as an effector of enhancer function.

To define the broader transcriptional consequences of H-ME perturbation, we performed RNA sequencing following CRISPRi-mediated enhancer repression in K562 cells. Differential expression analysis identified 187 significantly downregulated and 366 significantly upregulated genes, with *MYB* among the most strongly suppressed transcripts (Figure 3C). Gene set enrichment analysis revealed activation of differentiation-associated pathways, including heme metabolism, together with immune-related transcriptional programs (Figure 3D). Collectively, these findings demonstrate that H-ME maintains leukemic cell identity by sustaining *MYB* expression through an RNA-independent mechanism.

### Functional dissection identifies a GATA1/TAL1-dependent enhancer core within H-ME

Having established H-ME as a regulator of *MYB* expression, we next sought to define the functional sequences responsible for its activity. While CRISPRi identifies regulatory elements required for leukemic growth, it does not resolve the critical cis-regulatory sequences within an enhancer. We therefore performed high-resolution saturation mutagenesis using complementary CRISPR-Cas9 and CRISPRi tiling libraries spanning both the HBS1L-MYB-AHI1 locus and the H-ME region (Figure 4A-D).

**Figure 4:**
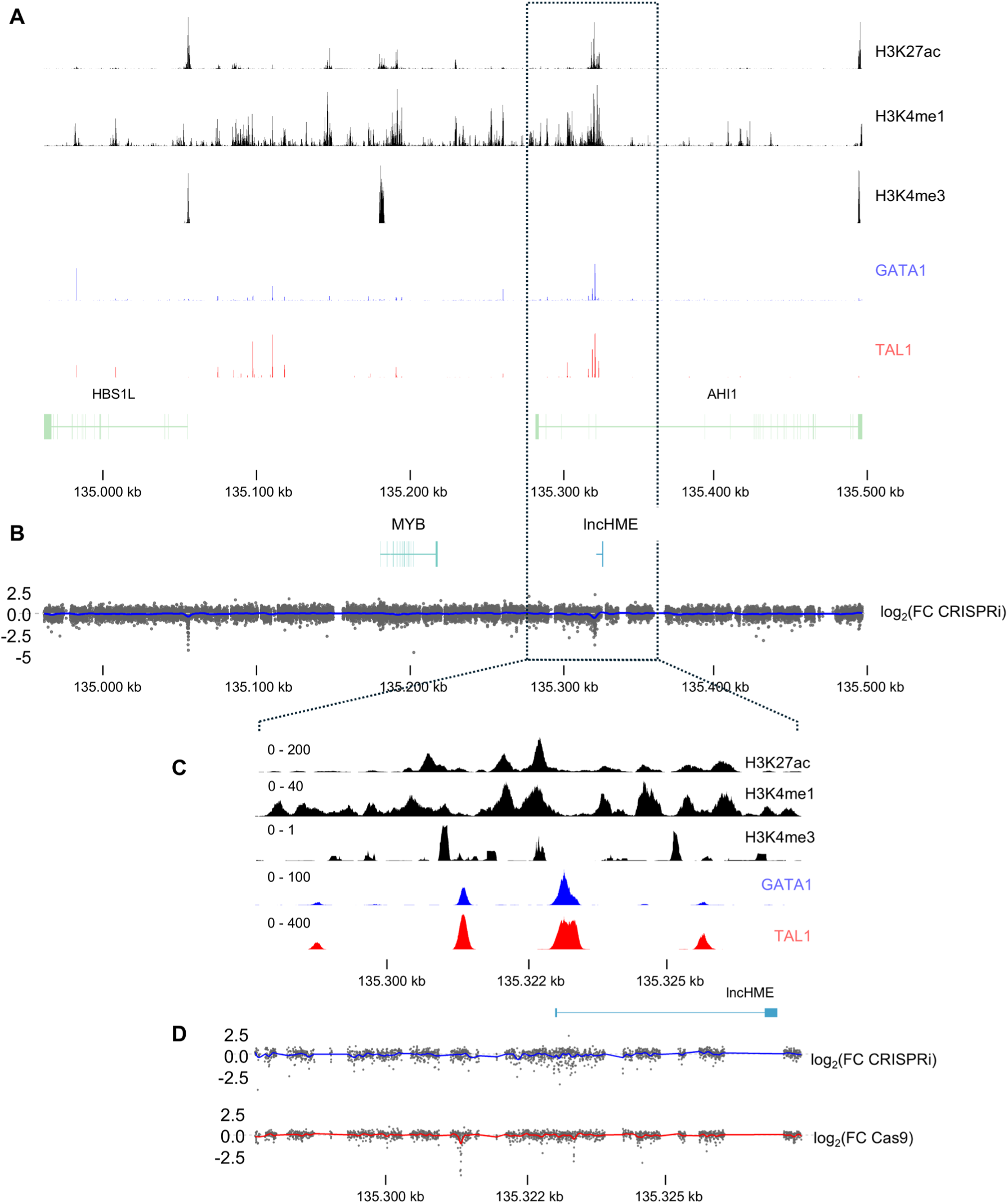
**A** and **C** Tracks from the publicly available ENCODE ChIP-Seq data in K562 showing from top to bottom: H3K27ac (black), H3K4me1 (black) and H3K4me3 (black), GATA1 (blue) and TAL1 (red) **B** and **D** Tiling screen results using CRISPRi with a smoothed fit curve (blue) and Cas9 with a smoothed fit curve (red) to analyze the essentiality of the HBS1L-MYB-AHI1 locus (n = 2 biological replicates) or the H-ME region (n = 2 biological replicates).

Analysis of sgRNA depletion identified a discrete ∼400 bp region within H-ME that accounted for the majority of the observed fitness phenotype. This essential enhancer core was independently identified by both the CRISPRi and Cas9 tiling screens, indicating that leukemic dependency is driven by a restricted set of regulatory sequences rather than the entire enhancer (Figure 4B,D).

To gain mechanistic insight into this functional enhancer core, we integrated the tiling screen results with publicly available ENCODE transcription factor ChIP-seq datasets (see Supplementary Table 5 for Details).^38^ The region of strongest functional dependency coincided with prominent binding sites for the hematopoietic transcription factors GATA1 and TAL1, both of which are key regulators of megakaryocytic differentiation and leukemic transcriptional programs (Figure 4A,C).^39–41^ Importantly, the sgRNAs producing the strongest depletion precisely overlapped these transcription factor-occupied regions, suggesting that GATA1/TAL1 binding is required for H-ME activity.

To validate these observations experimentally, we individually targeted the most strongly depleted regions identified in the saturation mutagenesis screen. Disruption of the GATA1/TAL1 binding motif significantly reduced expression of both lncHME and *MYB*, consistent with loss of enhancer activity (Supplementary Figure 4A-F). Likewise, CRISPR/Cas9-mediated disruption of either GATA1 or TAL1 reduced *MYB* expression (Supplementary Figure 4G).

To determine whether these transcription factors directly maintain enhancer activity, we performed ChIP-qPCR following transcription factor depletion. Targeting the TFBS within the H-ME enhancer led to loss of both GATA1 and TAL1 occupancy (Supplementary Figure 4H-I). Loss of either GATA1 or TAL1 resulted in decreased H3K27ac deposition at H-ME and at the MYB promotor (Supplementary Figure 4J-K). Together, these findings identify a GATA1/TAL1-dependent enhancer core that maintains H-ME activity and thereby sustains *MYB* expression in leukemic cells.

### H-ME is selectively active in megakaryocytic AML

Having identified a GATA1/TAL1-dependent enhancer core within H-ME, we next asked whether this regulatory circuit is active across all AML subtypes or restricted to specific leukemic lineages. We therefore analyzed chromatin accessibility in primary pediatric AML samples representing distinct genetic subgroups together with healthy hematopoietic progenitors.

ATAC-seq revealed markedly increased chromatin accessibility at both H-ME and the *MYB* promoter in non-Down syndrome acute megakaryoblastic leukemia (non-DS AMKL) and myeloid leukemia associated with Down syndrome (ML-DS), whereas accessibility was substantially lower in healthy CD34^+^ progenitors and in *KMT2A::MLLT3*-rearranged AML (Figure 5A-C). These findings indicate that H-ME is preferentially activated in megakaryocytic AML and suggest that enhancer accessibility is lineage restricted rather than a general feature of pediatric AML.

**Figure 5:**
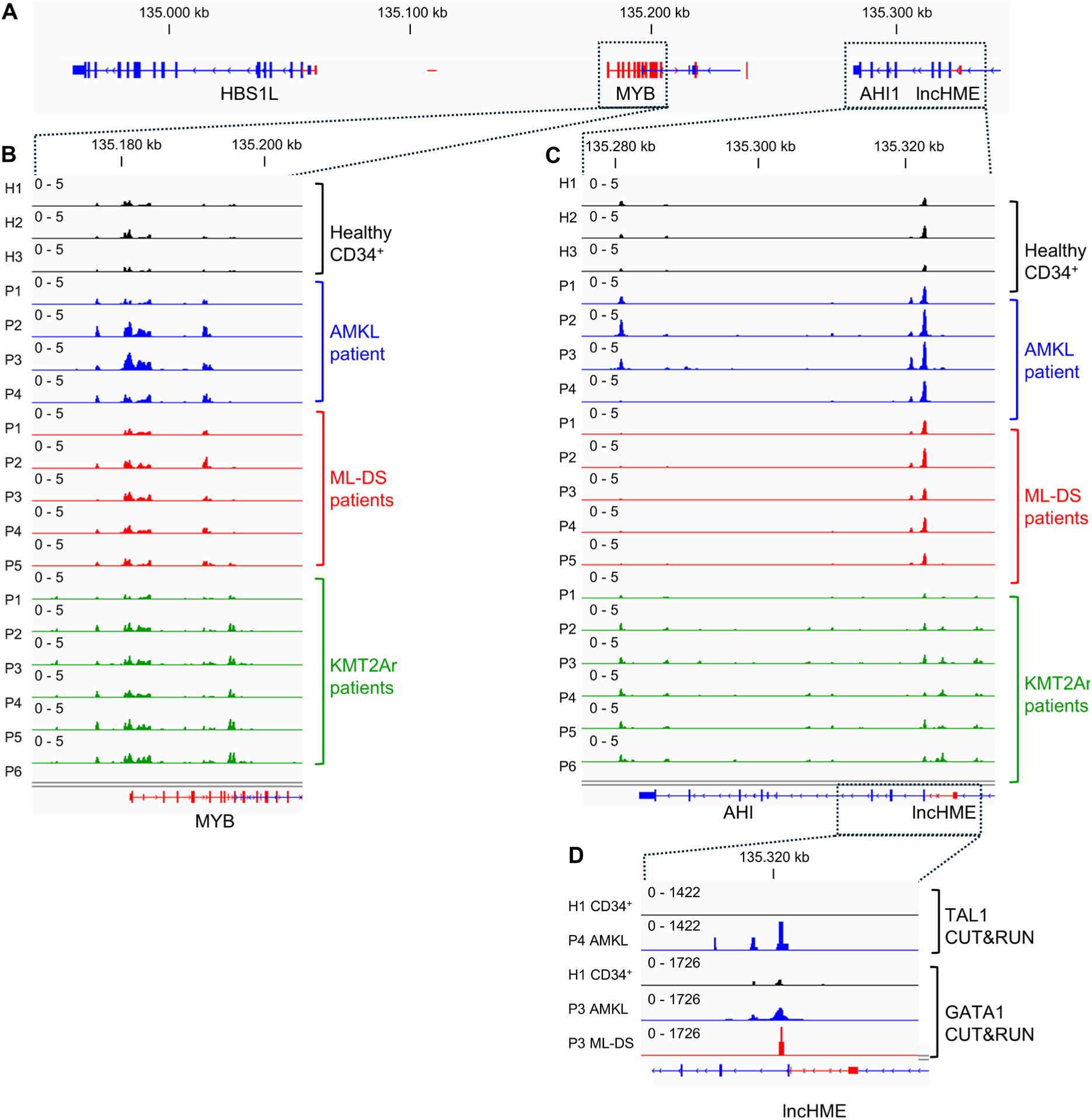
**A** IGV snapshot from the HBS1L-MYB-AHI locus on chromosome 6. **B** and **C** Tracks from ATAC-seq of CD34+ cells from healthy donors (black) and AML blasts from pediatric patients with different genetic backgrounds. Data from non-DS AMKL patients are shown in blue, ML-DS patients in red and KMT2A::MLLT3 patients in green. **D** Tracks from CUT&RUN experiments of GATA1 and TAL1 in pediatric patient samples with different genetic backgrounds. Non-DS AMKL patients are shown in blue, and ML-DS patients in red, compared with CD34+ cells from healthy donors (black).

To determine whether the transcription factor circuitry identified by saturation mutagenesis is maintained in primary leukemia, we performed CUT&RUN profiling for GATA1 and TAL1. Both transcription factors robustly occupied H-ME in primary non-DS AMKL and ML-DS samples, whereas binding was markedly reduced in healthy hematopoietic progenitors (Figure 5D and Supplementary Figure 5A-B). This occupancy closely mirrored the chromatin accessibility profiles and precisely overlapped the essential enhancer core identified by CRISPR tiling.

Together, these findings demonstrate that H-ME forms part of a lineage-specific regulatory circuit that is selectively engaged in megakaryocytic AML. The concordance between chromatin accessibility, transcription factor occupancy, and functional enhancer mapping strongly suggests that activation of H-ME is driven by a megakaryocytic GATA1/TAL1 transcriptional program that sustains *MYB* expression (Supplementary Figure 5C).

### H-ME is required for primary megakaryocytic AML growth in vitro and in vivo

Having established H-ME as a context-dependent regulator of *MYB*, we next asked whether this enhancer dependency is maintained in primary pediatric leukemia. To address this question, we investigated the functional consequences of H-ME perturbation in patient-derived models representing two genetically distinct megakaryocytic AML subtypes: non-Down syndrome acute megakaryoblastic leukemia (non-DS AMKL) and myeloid leukemia associated with Down syndrome (ML-DS).

Patient-derived leukemic cells expressing dCas9-KRAB were transduced with sgRNAs targeting H-ME and analyzed for clonogenic growth and gene expression. In non-DS AMKL-derived cells, CRISPRi-mediated repression of H-ME significantly reduced colony formation and was accompanied by decreased expression of both lncHME and *MYB* (Figure 6A-C). Similar results were observed in ML-DS-derived cells, where H-ME perturbation likewise impaired clonogenic growth and reduced *MYB* expression (Figure 6D-F). These findings demonstrate that H-ME dependency is conserved across genetically distinct megakaryocytic AML subtypes.

**Figure 6:**
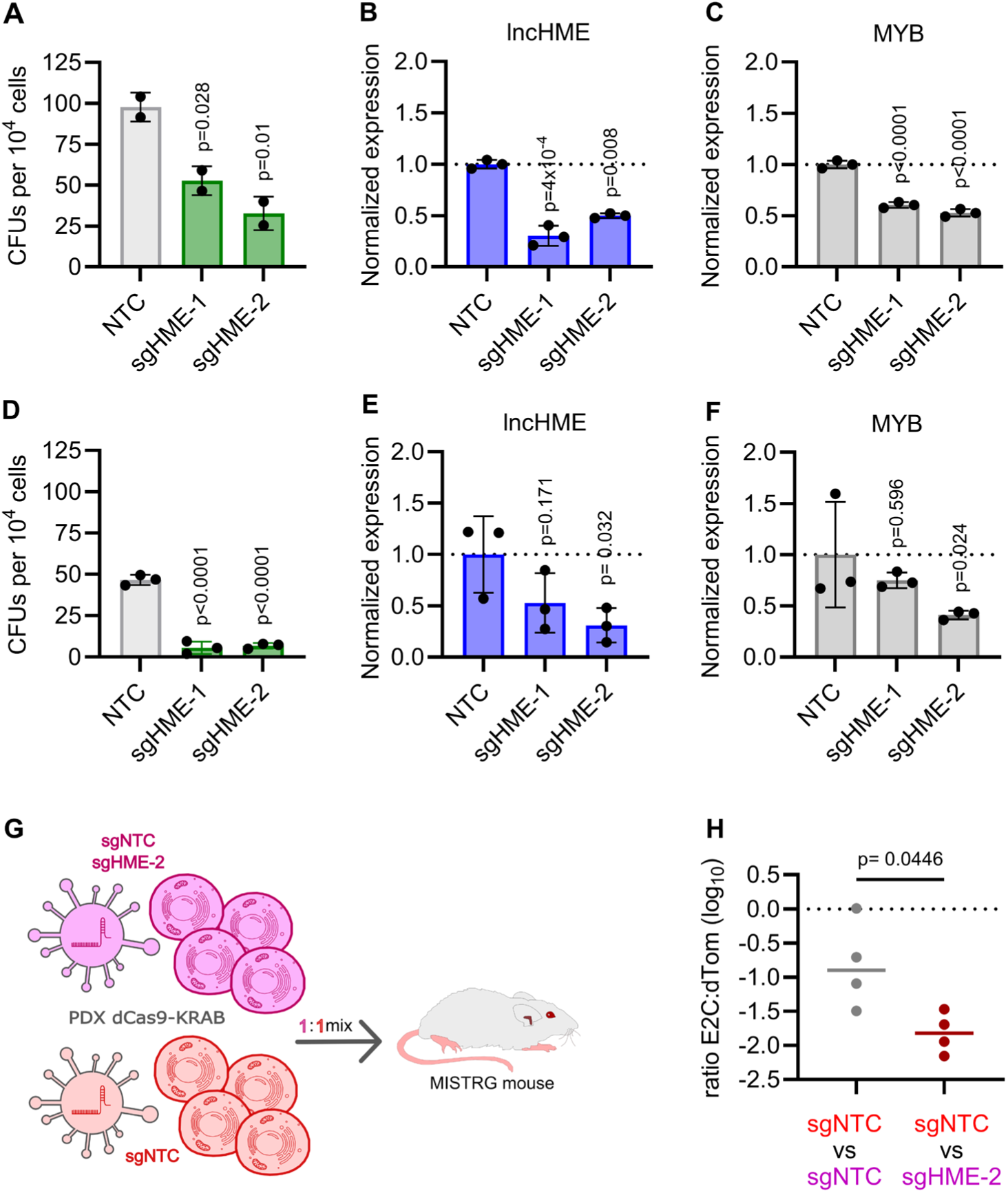
**A** Colony-forming assay following CRISPR knockdown of HME with two independent sgRNAs in non-DS AMKL PDX cells. Individual values are shown with the group means; n = 2 independent biological replicates. Statistics: one-way ANOVA on absolute colony counts (F(2,3) = 25.44; p = 0.0131) with Dunnett’s multiple comparisons test against the non-targeting control (NTC). **B** qPCR analysis of lncHME expression following CRISPR knockdown with two independent sgRNAs in non-DS AMKL PDX cells. Shown is the relative Expression (2^−ΔΔCt^), normalized to the housekeeping gene (B2M) and the non-targeting control (NTC); Error bars show the variance (asymmetric) back-transformed from the standard deviation of the ΔΔCt values, n = 3. Statistics: one-way ANOVA of the ΔCt values (F(2,6) = 30.86; p = 0.0007) with Dunnett post hoc test against NTC. **C** qPCR analysis of MYB expression following CRISPR knockdown with two independent sgRNAs in non-DS AMKL PDX cells. Shown is the relative Expression (2^−ΔΔCt^), normalized to the housekeeping gene (B2M) and the non-targeting control (NTC); Error bars show the variance (asymmetric) back-transformed from the standard deviation of the ΔΔCt values, n = 3. Statistics: one-way ANOVA of the ΔCt values (F(2,6) = 121.47; p = 0.0000) with Dunnett post hoc test against NTC. **D** Colony-forming assay following CRISPR knockdown of HME with two independent sgRNAs in ML-DS PDX cells. Individual values are shown with the group mean; n = 3 independent biological replicates. Statistics: one-way ANOVA on absolute colony counts (F(2,6) = 190.43; p = 0.0000) with Dunnett’s multiple comparisons test against the non-targeting control (NTC). **E** qPCR analysis of lncHME expression following CRISPR knockdown with two independent sgRNAs in ML-DS PDX cells. Shown is the relative Expression (2^−ΔΔCt^), normalized to the housekeeping gene (B2M) and the non-targeting control (NTC); Error bars show the variance (asymmetric) back-transformed from the standard deviation of the ΔΔCt values, n = 3. Statistics: one-way ANOVA of the ΔCt values (F(2,6) = 3.9; p = 0.0823) with Dunnett post hoc test against NTC. **F** qPCR analysis of MYB expression following CRISPR knockdown with two independent sgRNAs in ML-DS PDX cells. Shown is the relative Expression (2^−ΔΔCt^), normalized to the housekeeping gene (B2M) and the non-targeting control (NTC); Error bars show the variance (asymmetric) back-transformed from the standard deviation of the ΔΔCt values, n = 3. Statistics: one-way ANOVA of the ΔCt values (F(2,6) = 6.46; p = 0.0319) with Dunnett post hoc test against NTC. **G** Schematic diagram of the in vivo two-color competition assay using stably expressing dCas9-KRAB PDX cells. In brief, PDX cells are transduced with sgRNA virus for the non-targeting control (NTC) or the H-ME enhancer region. The cell populations are mixed in a 1:1 ratio (sgNTC vs sgNTC and sgNTC vs sgHME-2) and intravenously (i.v.) transplanted into immune-deficient MISTRG mice. **H** Competitive transplantation assay, endpoint analysis of bone marrow. Shown is the ratio of the APC⁺ population (sgHME or sgNTC) to the co-transplanted sgNTC-dTomato⁺ reference population within each mouse. Individual values with geometric mean, n = 4 mice per group. Statistics: unpaired t-test with Welch’s correction on ln-transformed ratios.

To determine whether this enhancer dependency is maintained under physiological conditions, we next performed competitive transplantation assays using ML-DS patient-derived xenografts (PDXs). Equal numbers of H-ME-targeted and control cells were differentially labeled, mixed at a 1:1 ratio, and transplanted into humanized immunodeficient mice (Figure 6G).^42^ Whereas both cell populations were equally represented at transplantation, H-ME-targeted cells were depleted at endpoint analysis, demonstrating a marked competitive disadvantage within the bone marrow compartment (Figure 6H and Supplementary Figure 6A).

Collectively, these findings establish H-ME as a bona fide enhancer dependency in primary pediatric megakaryocytic AML. The consistent effects observed across independent patient-derived models, distinct genetic backgrounds, and in vivo xenografts demonstrate that H-ME is not merely a cell line-specific regulator but a clinically relevant component of the leukemic transcriptional circuitry that sustains *MYB* expression and leukemic growth.

## Discussion

Epigenetic dysregulation is a defining feature of pediatric acute myeloid leukemia (pAML), yet identifying the cis-regulatory elements that actively maintain leukemic transcriptional programs remains a major challenge.^9^ While genome-wide epigenomic profiling readily identifies thousands of putative enhancers, only a minority represent functional dependencies required for leukemia maintenance. In this study, we developed an integrative strategy that combines H3K27ac profiling, enhancer-associated transcription, and functional CRISPRi screening to systematically prioritize biologically relevant enhancer elements. Using this approach, we validated H-ME as a lineage-restricted regulator of *MYB* that is required for leukemic growth. Mechanistically, H-ME maintains an active chromatin state at the *MYB* promoter, functions independently of its associated transcript, and is selectively active in megakaryocytic AML. Beyond the characterization of a single enhancer, our findings establish enhancer-associated transcription combined with CRISPRi screening as an effective strategy for systematic discovery of functional enhancer dependencies in pediatric AML.

A central conceptual advance of this work is the integration of enhancer-associated transcription with epigenomic profiling and CRISPRi screening. Although H3K27ac has become the standard marker for active enhancers, chromatin marks alone provide limited information regarding the functional importance of individual regulatory elements. Active enhancers frequently produce enhancer RNAs or longer non-coding transcripts, reflecting ongoing transcriptional activity at regulatory regions. We reasoned that combining enhancer-associated transcription with H3K27ac profiling would enrich for regulatory elements that are not only epigenetically active but also engaged within leukemic transcriptional networks. The successful validation of H-ME from more than 300 candidate enhancers demonstrates the utility of this strategy and suggests that integrating chromatin state with enhancer-associated transcription represents a broadly applicable framework for functional enhancer discovery in cancer.

*MYB* is one of the best-established oncogenic transcription factors in hematopoiesis and acute leukemia and is essential for maintaining proliferation while preventing differentiation. Multiple distal regulatory elements within the HBS1L-MYB locus have previously been implicated in controlling *MYB* expression during normal hematopoiesis and fetal hemoglobin regulation.^43,44^ More recently, chromatin looping and enhancer-associated regulatory mechanisms have been shown to contribute to *MYB* regulation in hematologic malignancies. During preparation of this manuscript, Mullin et al. identified the same enhancer, termed H-ME, and demonstrated its role in T-cell acute lymphoblastic leukemia through ETS1-and cBAF-dependent regulation. Complementary studies from the Hu laboratory further showed that H-ME regulates *MYB* expression through LDB1-mediated chromatin looping and ZNF217 in T-ALL. Our findings independently validate the biological importance of H-ME while substantially extending these observations. Specifically, we identify H-ME through an unbiased enhancer discovery pipeline, demonstrate that its biological activity is independent of the associated mature non-coding transcript, define a distinct GATA1/TAL1-dependent regulatory program in megakaryocytic AML, and establish its functional relevance in primary pediatric leukemia models. Together, these studies suggest that H-ME represents a conserved regulator of *MYB* whose upstream transcriptional circuitry is determined by cellular lineage.

Our study also contributes to the ongoing debate regarding the functional significance of enhancer-associated non-coding transcription. Although numerous enhancer-derived RNAs and long non-coding transcripts have been described whether these molecules actively mediate enhancer function or primarily reflect enhancer activity remains controversial. H-ME is associated with expression of lncHME, yet selective depletion of the transcript failed to alter *MYB* expression or leukemic proliferation. In contrast, perturbation of the underlying regulatory DNA element profoundly affected chromatin accessibility, enhancer activity, *MYB* expression, and leukemic fitness. These findings demonstrate that the biological activity of H-ME resides within the regulatory DNA element itself rather than its associated mature transcript. Collectively, these findings support the concept that enhancer-associated transcription can serve as a robust biomarker of functional enhancer activity, even when the transcript itself is dispensable.

An additional important finding is the preferential activity of H-ME in megakaryocytic AML. Chromatin accessibility, transcription factor occupancy, and functional perturbation consistently demonstrated preferential activation of H-ME in non-DS acute megakaryoblastic leukemia and myeloid leukemia associated with Down syndrome. The essential enhancer core identified by saturation mutagenesis coincided with binding of the megakaryocytic master regulators GATA1 and TAL1, indicating that H-ME functions within a lineage-specific transcription factor network that converges on *MYB*. This observation highlights how distinct hematopoietic transcriptional programs can utilize a common enhancer through different upstream regulatory mechanisms, thereby generating lineage-specific enhancer dependencies.

Finally, we demonstrate the biological relevance of H-ME in disease-relevant models. While many studies of enhancer biology rely predominantly on established cell lines, we validated H-ME function in primary patient-derived xenografts representing two genetically distinct megakaryocytic AML subtypes and confirmed its requirement for leukemic fitness in vivo. These findings indicate that enhancer dependency is maintained in primary leukemia and emphasize the importance of studying non-coding regulatory elements in clinically relevant experimental systems.

Collectively, our study establishes an integrative framework for the systematic identification of functional enhancer dependencies in pediatric AML. By combining enhancer-associated transcription, epigenomic profiling, and functional CRISPRi screening, we identify H-ME as an RNA-independent regulator of *MYB* that is essential for pediatric megakaryocytic leukemia. More broadly, our work demonstrates that enhancer-associated transcription provides an effective strategy for prioritizing functional regulatory elements and suggests that systematic interrogation of the non-coding genome can uncover previously unrecognized therapeutic vulnerabilities across hematologic malignancies.

## Acknowledgements

We thank F. Kalensee, V. Lang and K. Merkewitz (Goethe University Frankfurt) for technical assistance. We acknowledge E. Salzmann for statistical expertise and critical feedback on the analysis of experimental data. The study was supported by grants to JHK from the German Research Foundation (DFG; KL 2374/7-1) and Hilfe für Krebskranke Kinder e.V. as part of the C^3^OMBAT-AML research units. R.B. was a fellow of the Mildred Scheel Career Center (MSNZ). D.H. was supported by the Frankfurt Foundation for Children with Cancer and the Hilfe für Krebskranke Kinder e.V.

## Author contributions

L.S. performed experiments and analyzed the results. L.S., R.W., J. D. and K. S. performed the bioinformatic analysis and analyzed the results. L.V., H.I. and R.B. processed patient samples and generated data for the HemAtlas. R.B. and X. W. supervised the in vivo studies. R.C., Y.M. and M.K. designed and cloned the sgRNA libraries for the CRISPR screenings. S.H. contributed through funding acquisition for the research consortium RU5433 supporting this work. D.H. and J.H.K. designed and supervised the study. L.S., D.H., and J.H.K. crafted the manuscript. All authors revised and approved the manuscript.

## Data Availability Statement

Histone and transcription factor ChIP-seq data from ENCODE were used in this study (refer to Supplementary Table 5 for identifiers). The CRISPR screen, RNA-Seq, ATAC-Seq and CUT&Tag datasets generated during the current study have been deposited in NCBI’s Gene Expression Omnibus and are accessible through GEO Series accession number GSE343396-GSE43410. Respective sgRNA libraries are provided in with Supplementary Table 12 and Table 13 (Excel format). The datasets of pediatric AML patients analyzed during the current study have been published previously.^33,45^ This paper used existing analysis algorithms (see bioinformatic methods) and does not report original code. Any additional information required to reanalyze the data reported in this paper is available from the lead contacts upon request.

