## Supplementary Data for "Integrative enhancer discovery identifies functional enhancer dependencies in pediatric acute myeloid leukemia"

### Supplementary Figures

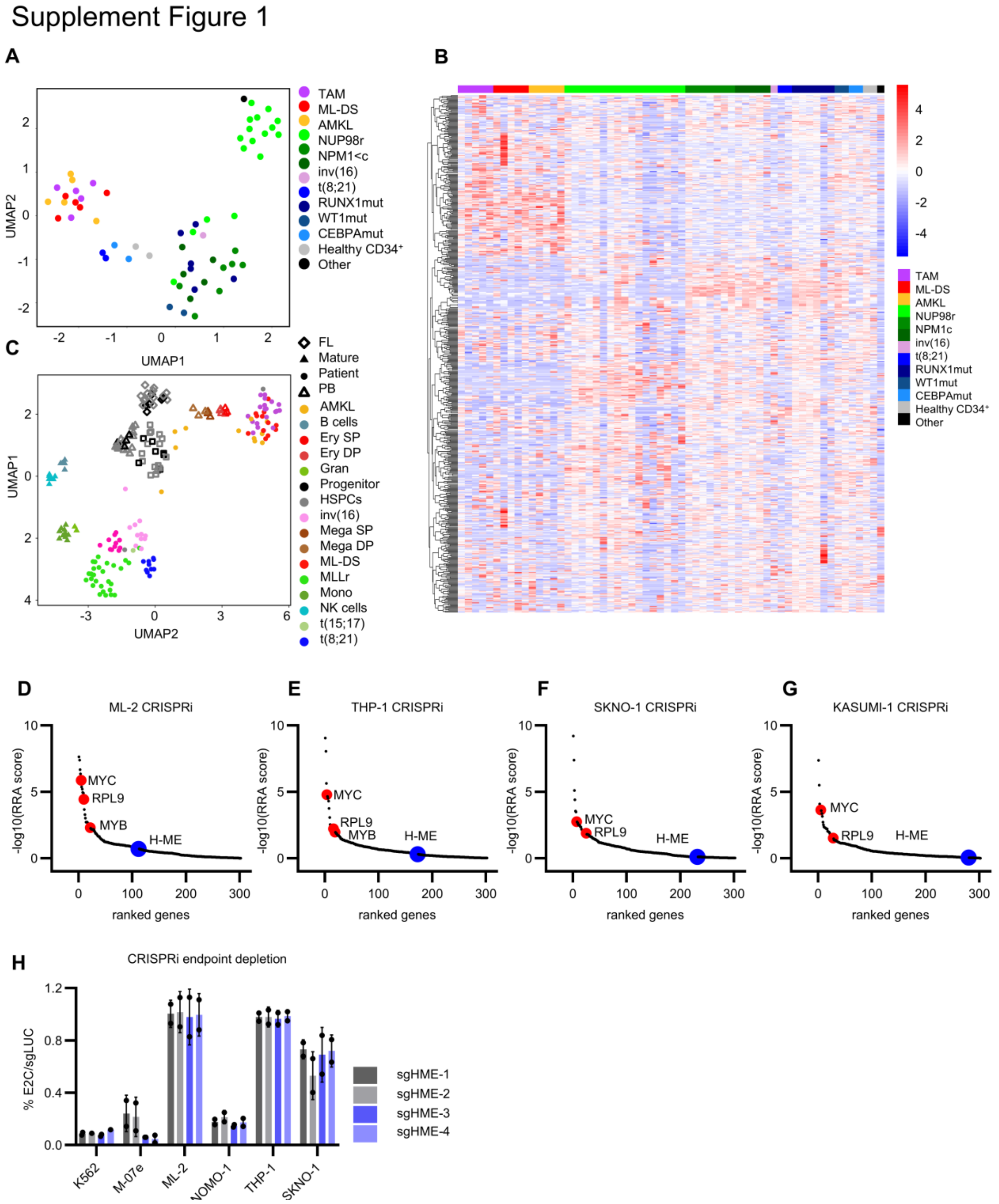


**Supplementary Figure 1: A** Dimensionality reduction of H3K27ac CUT&Tag data and a heatmap (**B**) from the hematopoietic database show clustering between healthy and leukemic samples. The database includes 83 patient samples, 13 healthy donor samples and 14 cell lines and PDX samples. Transient abnormal myelopoiesis (TAM), myeloid leukemia in Down syndrome (ML-DS), acute megakaryoblastic leukemia (AMKL).

**Supplementary Figure 1: (continued) C** Dimensionality reduction by UMAP of RNA-Seq data from the hematopoietic database shows clustering of long non-coding genes between healthy and leukemic samples. The database includes 96 healthy donor samples, 106 patient samples, and 22 PDX and cell lines. Fetal Liver (FL), peripheral blood (PB), acute megakaryoblastic leukemia (AMKL), single positive (SP), double positive (DP), Granulocytes (Gran), hematopoietic stem and progenitor cells (HSPCs), myeloid leukemia in Down syndrome (ML-DS), Monocytes (Mono), Natural Killer cells (NK). **D-G** Gene essentiality scores (RRA rank) from MAGeCK analysis of the CRISPRi screens in ML2, THP1, SKNO1 and Kasumi1 AML cell lines (n = 2 biological replicates each). The enhancer H-ME is highlighted in blue, and the positive controls RPL9, MYC, and MYB are highlighted in red. **H** Endpoint depletion values from fluorescence-based proliferation assays using CRISPRi H-Me MYB enhancer perturbation in different cell lines (n = 2 biological replicates, mean ± SD, 4 sgRNAs used for CRISPRi). The data are normalized to day 0 and to the non-targeting control (sgNTC).

Supplement Figure 2

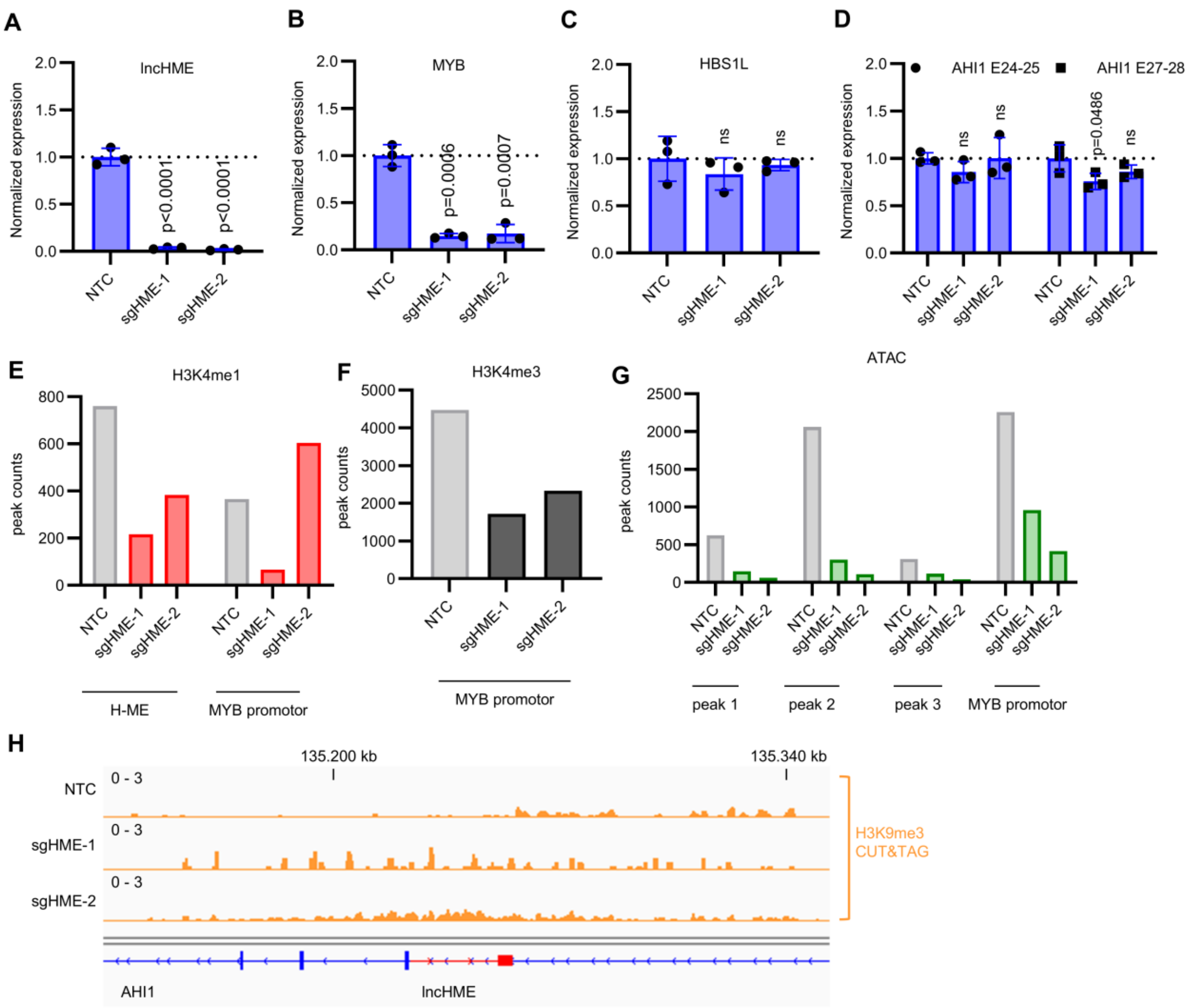


**Supplementary Figure 2: A** qPCR analysis of lncHME expression following CRISPR knockdown with two independent sgRNAs in K562 cells. Shown is the relative Expression (2^−ΔΔCt^), normalized to the housekeeping gene (B2M) and the non-targeting control (NTC); Error bars show the variance (asymmetric) back-transformed from the standard deviation of the ΔΔCt values, n = 3. Statistics: one-way ANOVA of the ΔCt values ($F(2,6) = 111.41$; $p = 0.0000$) with Dunnett post hoc test against NTC. **B** qPCR analysis of MYB expression following CRISPR knockdown with two independent sgRNAs in K562 cells. Shown is the relative Expression (2^−ΔΔCt^), normalized to the housekeeping gene (B2M) and the non-targeting control (NTC); Error bars show the variance (asymmetric) back-transformed from the standard deviation of the ΔΔCt values, n = 3. Statistics: one-way ANOVA of the ΔCt values ($F(2,6) = 35.35$; $p = 0.0005$) with Dunnett post hoc test against NTC. **C** qPCR analysis of HBS1L expression following CRISPR knockdown with two independent sgRNAs in K562 cells. Shown is the relative Expression (2^−ΔΔCt^), normalized to the housekeeping gene (B2M) and the non-targeting control (NTC); Error bars show the variance (asymmetric) back-transformed from the standard deviation of the ΔΔCt values, n = 3. Statistics: one-way ANOVA of the ΔCt values ($F(2,6) = 0.59$; $p = 0.5852$) with Dunnett post hoc test against NTC. **D** qPCR analysis of AHI1 expression following CRISPR knockdown with two independent sgRNAs in K562 cells. Shown is the relative Expression (2^−ΔΔCt^), normalized to the housekeeping gene (B2M) and the non-targeting control (NTC); Error bars show the variance (asymmetric) back-transformed from the standard deviation of the ΔΔCt values, n = 3. Statistics: one-way ANOVA of the ΔCt values ($F(2,6) = 1.17$; $p = 0.3722$, $F(2,6) = 4.17$; $p = 0.0734$) with Dunnett post hoc test against NTC. **E-G** Quantified CUT&Tag data (peak counts) for H3K4me1 (red), H3K4me3 (black), and ATAC (green) for H-ME region and the MYB promoter upon CRISPRi perturbation. **H** Tracks from the CUT&Tag of H3K9me3 experiment upon CRISPRi targeting H-ME region.

Supplement Figure 3

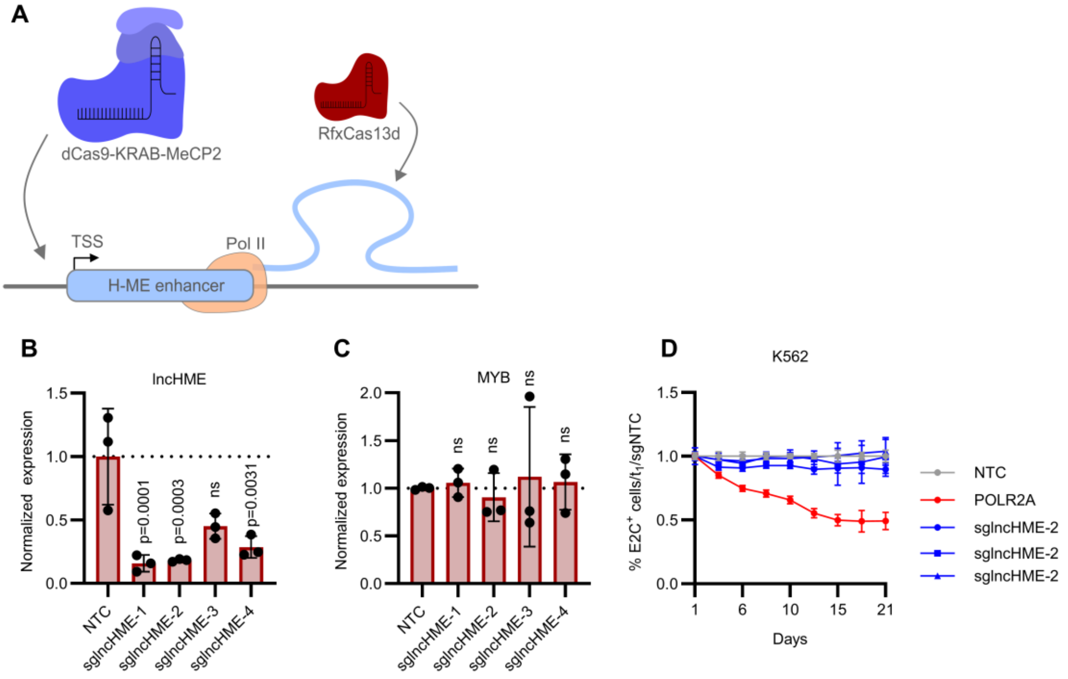


**Supplementary Figure 3: A** Schematic representation of the CRISPRi system compared to the RfxCas13d (CasRX) system. **B** qPCR analysis of lncHME expression following CasRX knockdown with four independent sgRNAs in K562 cells. Shown is the relative Expression (2^−ΔΔCt^), normalized to the housekeeping gene (B2M) and the non-targeting control (NTC); Error bars show the variance (asymmetric) back-transformed from the standard deviation of the ΔΔCt values, n = 3. Statistics: one-way ANOVA of the ΔCt values ($F(4,10) = 15.9$; $p = 0.0002$) with Dunnett post hoc test against NTC. **C** qPCR analysis of MYB expression following CasRX knockdown with four independent sgRNAs in K562 cells. Shown is the relative Expression (2^−ΔΔCt^), normalized to the housekeeping gene (B2M) and the non-targeting control (NTC); Error bars show the variance (asymmetric) back-transformed from the standard deviation of the ΔΔCt values, n = 3. Statistics: one-way ANOVA of the ΔCt values ($F(4,10) = 0.13$; $p = 0.9691$) with Dunnett post hoc test against NTC. **D** Proliferation assay of K562 cells upon CRISPR RNA-targeting of the lncHME transcript. The different sgRNAs are shown in blue, the positive control sgRNAs for POLR2A is shown in red and the non-targeting control is shown in grey.

#### Supplement Figure 4

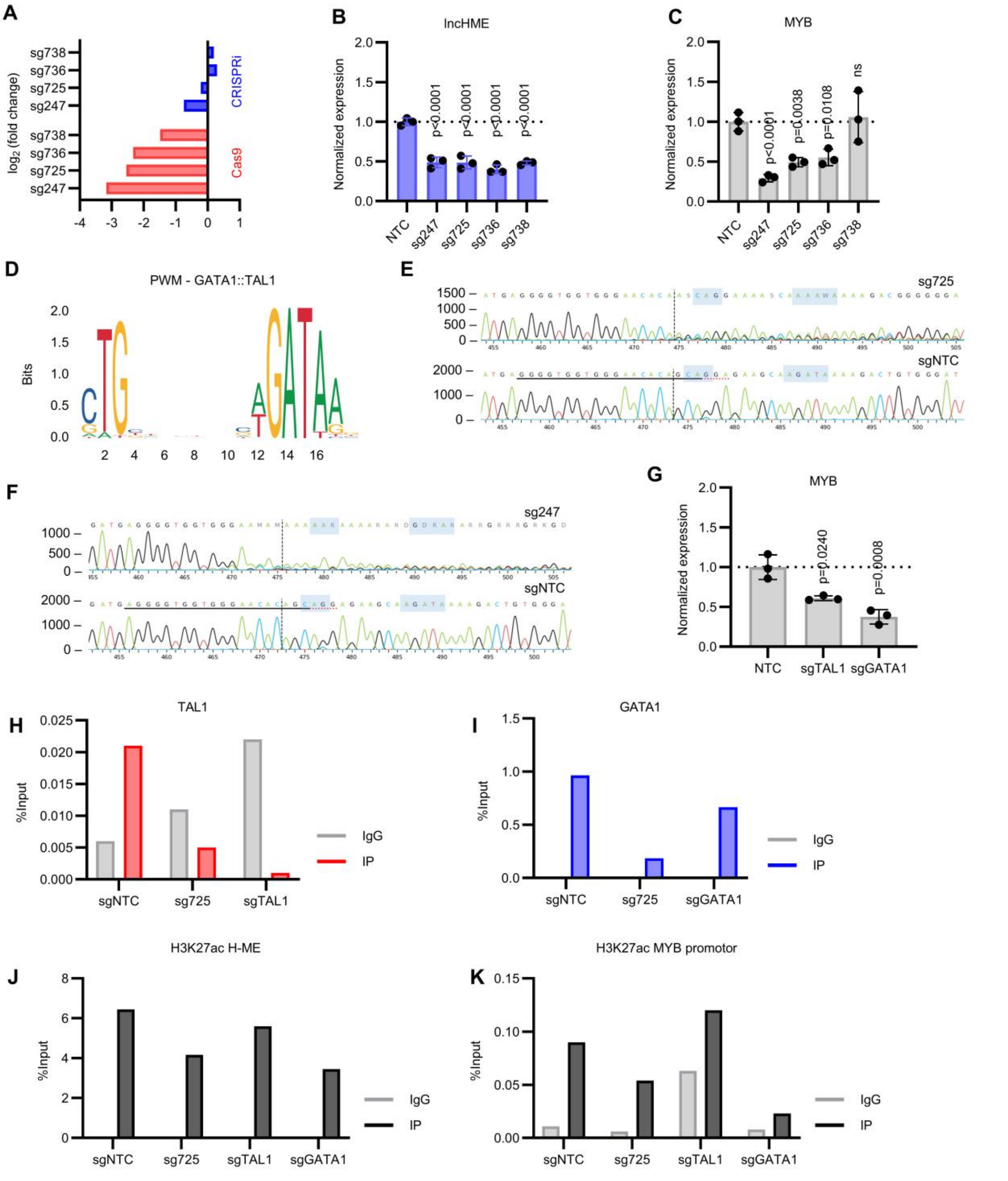


**Supplementary Figure 4: A** Log2 fold change of most depleted sgRNAs from the tiling screen with Cas9 (red) and CRISPRi (blue). **B** qPCR analysis of lncHME expression following CRISPR knockout with four independent sgRNAs in K562 cells. Shown is the relative Expression ($2^{-\Delta\Delta Ct}$), normalized to the housekeeping gene (B2M) and the non-targeting control (NTC); Error bars show the variance (asymmetric) back-transformed from the standard deviation of the $\Delta\Delta Ct$ values, n = 3. Statistics: one-way ANOVA of the $\Delta Ct$ values ($F(4,10) = 27.4$; $p = 0.0000$) with Dunnett post hoc test against NTC. **C** qPCR analysis of MYB expression following CRISPR knockout with four independent sgRNAs in K562 cells. Shown is the relative Expression ($2^{-\Delta\Delta Ct}$), normalized to the housekeeping gene (B2M) and the non-targeting control (NTC); Error bars show the variance (asymmetric) back-transformed from the standard deviation of the $\Delta\Delta Ct$ values, n = 3. Statistics: one-way ANOVA of the $\Delta Ct$ values ($F(4,10) = 22.83$; $p = 0.0001$) with Dunnett post hoc test against NTC.

**Supplementary Figure 4: (continued) D** Position weight matrix of the GATA1::TAL1 (MA0140.2) from the JASPAR database. **E** Tracking of Indels by Decomposition (TIDE) analysis of sg725 from the saturation mutagenesis screening. In blue, the GATA1::TAL1 motif is highlighted. Lower sequences show the non-targeting control. **F** Tracking of Indels by Decomposition (TIDE) analysis of sg247 from the saturation mutagenesis screening. In blue, the GATA1::TAL1 motif is highlighted. Lower sequences show the non-targeting control. **G** qPCR analysis of MYB expression following CRISPR knockout of TAL1 and GATA1 with one independent sgRNAs each in K562 cells. Shown is the relative Expression ($2^{-\Delta\Delta Ct}$), normalized to the housekeeping gene (B2M) and the non-targeting control (NTC); Error bars show the variance (asymmetric) back-transformed from the standard deviation of the ΔΔCt values, n = 3. Statistics: one-way ANOVA of the ΔCt values ($F(2,6) = 24.54$; $p = 0.0013$) with Dunnett post hoc test against NTC. **H** ChIP-qPCR of TAL1 occupancy at H-ME upon TFBS disruption with sg725 and TAL1 CRISPR/Cas9 knockout. **I** ChIP-qPCR of occupancy at the MYB promoter upon TFBS disruption with sg725 and GATA1 CRISPR/Cas9 knockout. **J** ChIP-qPCR of H3K27ac at H-ME upon TFBS disruption with sg725 and GATA1 and TAL1 CRISPR/Cas9 knockout. **K** ChIP-qPCR of H3K27ac at the MYB promoter upon TFBS disruption with sg725 and GATA1 and TAL1 CRISPR/Cas9 knockout.

Supplement Figure 5

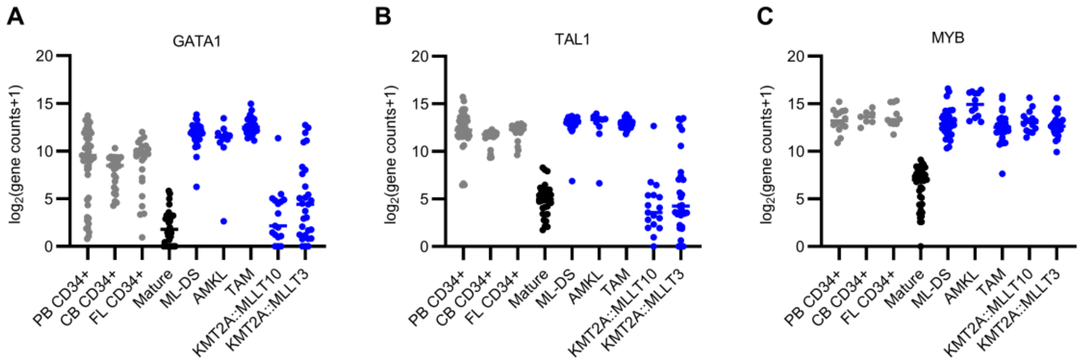


**Supplementary Figure 5: A** GATA1 expression from our RNA-seq hematopoietic database (HemAtlas) in healthy progenitor cells (grey), mature cells (black) and different pediatric AML subtypes (blue). **B** TAL1 expression from our RNA-seq hematopoietic database (HemAtlas) in healthy progenitor cells (grey), mature cells (black) and different pediatric AML subtypes (blue). **C** MYB expression from our RNA-seq hematopoietic database (HemAtlas) in progenitor cells (grey), mature cells (black) and different pediatric AML subtypes (blue).

### Supplement Figure 6

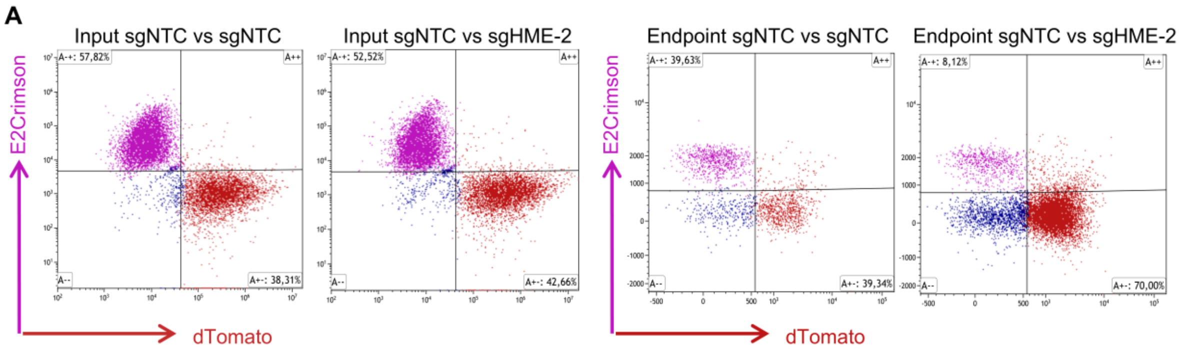


**Supplementary Figure 6: A** Representative input (left) and endpoint (right) flow cytometry data from the two-color competitive experiment.

### Supplementary Materials and Methods

#### Library Screenings

Screenings were conducted in seven AML cell lines (K562, M-07e, NOMO-1, THP-1, ML-2, KASUMI-1, SKNO-1). Specifically, AML cell lines were transduced with lentivirus carrying the sgRNA library (MOI 0.3) and maintained in culture until they reached at least 14 doublings. Cells were collected initially and at mid and late time points, which were determined by the depletion pattern of essential genes like POLR2A or RPL9. Then, genomic DNA was isolated from the cell lines, and the sgRNA sequences were amplified using PCR with the 2x NEB Next Master Mix (#M0541, New England Biolabs, Frankfurt am Main, Germany). Next-generation sequencing (Illumina NovaSeq, PE150) of all samples was performed and analyzed by using the MaGeCK algorithm.[1] Target genes were selected based on their fold change and significance. Additional filtering was performed using essentiality scores from the DepMap (https://depmap.org/portal/) and relevant publications.[2]

#### Cell culture

AML cell lines were obtained from the Leibniz Institute DSMZ-German Collection of Microorganisms and Cell Cultures GmbH (DSMZ, Braunschweig, Germany, see Supplementary Table 6 for Details). Cell lines were maintained in RPMI 1640 GlutaMAX™ (Cat#61870036, Thermo Fisher Scientific, Darmstadt, Germany) supplemented with 10% or 20% fetal calf serum (FCS), 1% Penicillin-Streptomycin (Thermo Fisher Scientific, Cat#15070063) and additional human cytokines if needed (see Supplementary Table 7). Cytokines were obtained from Thermo Fisher Scientific. HEK293T cells were obtained from DSMZ and cultured in DMEM (Thermo Fisher Scientific, Cat#21969035) supplemented with 10% FCS and 1% Penicillin-

Streptomycin. All cell lines were routinely tested for mycoplasma contamination. Cell line characteristics, including sex and age, can be found in Supplementary Table 8.

Hematopoietic primary cells and patient-derived xenograft cells were cultured in Stemspan (Cat#09650, Stem Cell Technologies, Köln, Germany) supplemented with 50 ng/ml human SCF, 50 ng/ml human FLT3-L, 10 ng/ml human IL-6, 2.5 ng/ml human IL-3, and 20 ng/ml human TPO and 1% Penicillin-Streptomycin. Stable dCas9-KRAB and Cas9 cell lines and PDXs were established via lentiviral transduction (Addgene #90332, #57821, or #170482).[3–5]

#### Transduction and hematopoietic assays

CRISPR cell lines were thawed and selected with Blasticidin for at least seven days before transduction. For gene expression and epigenetic profiling experiments, CRISPRi cells were sorted five days after sgRNA transduction, and Cas9 cells were sorted three days after transduction.

Patient-derived xenograft cells expressing the dCas9-KRAB protein were generated by lentiviral transduction and serially transplanted into immunodeficient MISTRG mice (see Supplementary Table 6 and 9 for Details). For colony formation assays and qRT-PCR experiments, PDX cells were isolated from MISTRG mice and recovered for at least 48 hours. PDX cells were transduced with sgRNA virus and sorted two days after transduction. For colony formation assays, 10 000 cells were seeded per dish and counted after 10 days.

#### Flow cytometry and cell sorting

Flow cytometry analysis of PDX cells and AML cell lines was performed using a CytoFLEX S (Beckman Coulter, Krefeld, Germany) device. Cell lines were sorted with the CytoFLEX SRT Benchtop Cell Sorter (Beckman Coulter), and PDX cells were sorted using a FACSAria™ III Cell Sorter (BD Bioscience, Heidelberg, Germany).

#### qRT-PCR and RNA-Seq

Cell lines and PDX cells were transduced with sgRNA virus (see Supplementary Table 10), then harvested and sorted two or three days after transduction. RNA was isolated using the Quick-RNA Microprep Kit (Cat#R1050, Zymo Research, Freiburg, Germany).

For gene expression analysis with qRT-PCR, cDNA was synthesized using the High-Capacity cDNA Reverse Transcription Kit (Thermo Fisher Scientific, Cat#4374967). qRT-PCR was performed with forward and reverse exon-spanning primers (Supplementary Table 11) and SYBR™ Select Master Mix (Thermo Fisher Scientific, Cat#4472919) on a QuantStudio™ 5 cycler (Thermo Fisher Scientific).

RNA sequencing was performed by Novogene Company, Ltd. (München, Germany) using rRNA-depleted RNA on an Illumina NovaSeq X using 150 bp paired-end sequencing. The raw sequencing data were processed using the nf-core/rnaseq pipeline[6,7] with default settings as previously described in Winkler et al.[8] Cutting efficacy of all CRISPR/Cas9 sgRNAs used was determined in advance of the experiment as described in Verboon et al.[9]

#### Detailed Statistical Analysis

For the Rescue assay fractions of transduced cells were determined by flow cytometry at days 1, 6 and 11. For each culture, values were normalized to the corresponding day 1 measurement. Group differences in the extent of decline were assessed by two-way ANOVA (group × time) with repeated measures across timepoints, followed by Dunnett's multiple comparisons test against the non-targeting control at each timepoint

(GraphPad Prism 11, GraphPad Software, Boston, MA, USA, www.graphpad.com) Day 1 was excluded from post-hoc testing as normalization constrains it to unity in all groups. n = 2 independent transductions per group; multiplicity-adjusted two-sided p-values $< 0.05$ were considered significant.

Colony-forming capacity was assessed in methylcellulose, and colonies were counted at endpoint. Statistical testing was performed on absolute colony counts rather than on control-normalized values, as normalization to the control mean removes the variance of the control group from the error estimate. Group differences were assessed by one-way ANOVA followed by Dunnett's multiple comparisons test against the non-targeting control (GraphPad Prism 11). Sample sizes are indicated in the figure legends; multiplicity-adjusted two-sided p values $< 0.05$ were considered significant.

Relative gene expression was quantified by the $\Delta\Delta Ct$ method. All statistical testing was performed on $\Delta Ct$ values rather than on fold-change values, since $\Delta Ct$ is approximately normally distributed with homogeneous variance whereas $2^{-\Delta Ct}$ values are log-normally distributed. Group differences were assessed by one-way ANOVA followed by Dunnett's multiple comparisons test against the non-targeting control (GraphPad Prism 11). Fold-change values are shown for visualization only, with error bars derived by back-transformation of the mean ± SD of $\Delta\Delta Ct$. n = 3 per group; adjusted two-sided p values $< 0.05$ were considered significant.

Bone marrow was analyzed at endpoint by flow cytometry. For each mouse, we calculated the ratio of the $APC^+$ population (sgHME or sgNTC) to the co-transplanted sgNTC-$dTomato^+$ reference population, providing an internal control for engraftment efficiency. Ratios were ln-transformed prior to analysis, as ratio data are right-skewed and bounded at zero; group means were compared by unpaired t-test with Welch's correction (GraphPad Prism v11). Group summary values are reported as geometric means. n = 4 mice per group; two-sided p values < 0.05 were considered significant.

### Supplementary Tables

**Supplementary Table 1:** Patient information non-DS AMKL and ML-DS

| Figure ID | **AMKL P1** | **AMKL P2** | **AMKL P3** | **AMKL P4** | **ML-DS P1** | **ML-DS P2** | **ML-DS P3** | **ML-DS P4** | **ML-DS P5** |
|---|---|---|---|---|---|---|---|---|---|
| sex | f | f | m | m | m | m | m | m | f |
| age at diagnosis | 1y 4m | 0y 1m | 0y 6m | 0y 3m | 2y 6m | 1y 6m | 2y 4m | 1y 9m | 1y 11 m |
| WBC ($x10^9$/L) | 33.05 | 26.61 | 32.05 | 106.46 | 4.35 | 22.79 | 3.89 | 3.8 | 168 |
| Hb (g/dl) | 11.7 | 8.6 | 6.4 | 7.9 | 7.6 | 5.4 | 7.8 | 8.7 | 7.1 |
| BM blasts (%) | 34 | 17 | 56 | 74 | 61.5 | 75.5 | 2 (PB) | 1 | 73 |
| CNS | No | N/A | No | No | No | No | No | No | No |
| SCT | No | Yes | Yes | No | No | No | Yes | Yes | No |
| molecular mutations | JAK2 | N/A | N/A | N/A | | IRX1, JAK1, MBNL1, NRAS | EZH2 | JAK3, KMT2C | NRAS |
| Cytogenetics (karyotype) | 46, XX, add(11)(p15), add(12)(p13) ,del(13)(q14q21),del(17)(q22)(18)/46,XX[7] | 46,XX[15].nuc ish 11q23(MLLx2)[100/100], nuc ish 3q26(EVI1x2)[100/100],8q22(RUNX1T1 x2),21q22(RUNX1x2) [100/100],11q23(MLLx2)[100/100], 16q22(CBFBx2)[100/100], 17q21.1(RARAx2)[100/100] | 48,XY,+6,+6[1]/48,idem,del(3)(q13q26),add(11) (p14)[23]/46,XY[1] | 46,XY[15].nuc ish 3q26(EVI1x2)[100/100],8q22(RUNX1T1x2) ,21q22(RUNX1x2)[100/100], 11q23(MLLx2)[100/100], 6q22(CBFBx2)[99/100], 17q21.1(RARAx2)[100/100] | N/A | no abberations | trisomy 8 | N/A | N/A |
| response | PR nach HAM | PR: nach AI, CCR: haM | CCR: nach HAM | CCR: nach HAM | CRp nach AIE | CRp nach 1. CPX-351 | CRp nach 1. CPX-351 | CRp nach 1. CPX-351 | CCR |
| relapse | Yes | Yes | No | No | No | No | Yes | No | No |

**Supplementary Table 2:** Patient information KMT2A::MLLT3

| **Figure ID** | **KMT2A:: MLLT3 P1** | **KMT2A:: MLLT3 P2** | **KMT2A:: MLLT3 P3** | **KMT2A:: MLLT3 P4** | **KMT2A:: MLLT3 P5** | **KMT2A:: MLLT3 P6** |
|---|---|---|---|---|---|---|
| sex | f | f | f | m | m | f |
| age at diagnosis | 12y 10m | 17y 6m | 15y 1m | 0y 5m | 15y 11m | 6y 2m |
| WBC (x$10^9$/L) | 1.09 | 9.6 | 119.03 | 103.85 | 8.6 | N/A |
| Hb (g/dl) | 8.3 | 6.5 | 10.6 | 7.3 | 7.2 | N/A |
| BM blasts (%) | 77 | 68 | 91 | 86 | 55 | N/A |
| CNS | No | No | No | No | No | N/A |
| SCT | No | No | No | No | No | No |
| molecular mutations | FLT3, U2AF1 | NRAS | N/A | N/A | N/A | FLT3_TKD, ACVR1B, ATIC, CD79B, FANCM, MSH3, PER1, TPBG |
| Cytogenetics (karyotype) | 49,XX,+i(8)(q10),t(9;11)(p21;q23),+19,+mar[4]/48,idem,i(8)(q10),-mar[11] | 46,XX,t(9;11)(p21;q23)[12]/47,idem,+8[3] | 47,XX,+dmin[13]/46,XX[2].nuc ish 3q26(EVI1x2)[100/100],8q22(RUNX1T1 x2),21 q22(RUNX1x2)[100/100], 11q23(MLLx3)[92/100], 16q22(CBFBx2)[100/100],17q21.1(RARAx2)[100/100] | 46,XY, t(9;11)(p21;q23)[16].nuc ish 3q26(EVI 1x2)[100/100],8q22(RUNX1T1x2),21 q22(RUNX1x2)[99/100],9q34(ABL 1x2),22q 11(BCRx2)[98/100],11 q23(MLLx2)(5' MLL sep 3'MLLx 1)[92/100],12p13(ETV6x2),21q22(RUNX 1x2)[97/100], 16q22(CBFBx2)[100/100], 17q21.1(RARAx2)[98/100] | 46,XY, t(9; 11 )(p21 ;q23)[1]/47, idem,+8[19]/48,idem,+3,+8[2].nuc ish 3q26(EVI 1x2)[100/100],8q22(RUNX1T1x3),21q22(RUNX1x2)[95/100], 16q22(CBFBx2)[100/100], 17q21.1(RARAx2)[100/100] | 46,XX,del(9)(p13p21),t(9;11)(p22;q23),t(13;17)(q21;q24)[15] |
| response | CCR nac HAM | PR: nach HAM; CCR: nach AI | CCR: nach HAM | CCR: nach HAM | CCR: nach HAM | CCR |
| relapse | No | No | No | No | No | No |

**Supplementary Table 3: Information about Antibodies used in CUT&Tag and ChIP**

| Antibodies | Source | Identifier |
|---|---|---|
| Anti-humanCD45-APC/Cyanine7 | Biolegend | Cat#368516 |
| H3K4me3 antibody used for CUT&TAG | Epicypher | Cat#13-0060 |
| H3K4me1 antibody used for CUT&TAG | Epicypher | Cat#13-0040 |
| H3K27ac antibody used for CUT&TAG and ChIP | Diagenode | Cat#15410174 |
| H3K9me3 antibody used for CUT&TAG | Abcam | Cat#8898 |
| TAL1 antibody used for ChIP | cell signaling | Cat#12831S |
| GATA1 antibody used for ChIP | abcam | Cat#11852 |
| rabbit IgG antibody used for ChIP | diagenode | Cat#C15410206 |

**Supplementary Table 4:** ChIP qPCR primer

| Name | Primer sequence |
|---|---|
| FW_enhancer_ChIP | TGTAGTCATAAAGAGCCACTACCT |
| RV_enhancer_ChIP | GCAAGAGGAAGTGAATGTTGGG |
| FW_MYB_promotor_ChIP | GCACAGTTGTAAACCTTGACG |
| RV_MYB_promotor_ChIP | TCCAGCTCCCACTCACT |
| FW_H3K27ac_ChIP | GCAGTTGGGGAACTAGAAGCT |
| RV_H3K27ac_ChIP | GCCTCATGCACAGTTCTGTC |

**Supplementary Table 5:** Information for ENCODE identifiers

| Other | Source | Identifier |
|---|---|---|
| K562 ChIP-seqdata: H3K4Me3 | ENCODE | ENCFF144MRB |
| K562 ChIP-seqdata: H3K4Me1 | ENCODE | ENCFF100FDI |
| K562 ChIP-seqdata: H3K27Ac | ENCODE | ENCFF094XCU |
| K562 ChIP-seqdata: TAL1 | ENCODE | ENCFF700NBW |
| K562 ChIP-seqdata: GATA1 | ENCODE | ENCFF331URE |

**Supplementary Table 6.** Information for experimental models

| Cell lines | Source | Identifier |
|---|---|---|
| HEK293T | DSMZ | DSMZ#ACC635; RRID:CVCL_0063 |
| K562 | DSMZ | DSMZ#ACC10; RRID:CVCL_0004 |
| M-07E | DSMZ | DSMZ#ACC104; RRID:CVCL_2106 |
| NOMO-1 | DSMZ | DSMZ#ACC542;RRID:CVCL_1609 |
| THP1 | DSMZ | DSMZ#ACC16;RRID:CVCL_0006 |
| ML-2 | DSMZ | DSMZ#ACC15; RRID:CVCL_1418 |
| KASUMI-1 | DSMZ | DSMZ#ACC220; RRID:CVCL_0589 |
| SKNO-1 | DSMZ | DSMZ#ACC690; RRID:CVCL_2196 |
| **Mouse strains** | **Source** | **Identifier** |
| Mus musculus: MISTRG (Rag2ko/ko, Csf2/Il3ki/ki, Il2rgko/ko, Thpoki/ki, hSIRPA+/+, Csf1ki/ki) | Regeneron Pharmaceuticals | not commercially available |

**Supplementary Table 7:** Information about human cytokines

| Chemicals, peptides and recombinant proteins | Source | Identifier |
|---|---|---|
| Recombinant human SCF | Thermo Fisher | Cat#300-07 |
| Recombinant human FLT3L | Thermo Fisher | Cat#300-19 |
| Recombinant human IL-3 | Thermo Fisher | Cat#200-03 |
| Recombinant human IL-6 | Thermo Fisher | Cat#200-06 |
| Recombinant human TPO | Thermo Fisher | Cat#300-18 |
| Recombinant human GM-CSF | Thermo Fisher | Cat#300-03 |
| Stem Regenin 1 | STEMCELL | Cat#72344 |
| UM171 | STEMCELL | Cat#72914 |

**Supplementary Table 8:** Cell line information

| Cell line | K562 | M-07E | NOMO-1 | THP-1 | ML-2 | KASUMI-1 | SKNO-1 |
|---|---|---|---|---|---|---|---|
| Adult (A)/Pediatric (P) | A | P | A | P | A | P | A |
| Origin age (years) | 53 | 0.5 | 31 | 1 | 26 | 7 | 22 |
| Origin sex | f | f | f | m | m | m | m |
| Leukemia classification | CML | AML M7 | AML M5a | AML M5a | AML M4 | AML M2 | AML M2 |
| Fusion protein | BCR::ABL | | MLL:: MLLT3 | MLL:: MLLT3 | MLL::MLLT4 | RUNX1:: RUNX1T1 | RUNX1:: RUNX1T 1 |
| Specific translocation | t(9;22)(q34;q11) | | t(9;11)(p22;q23) | t(9;11)(p21;q23) | t(6;11)(q27;q23) | t(8;21)(q22;22) | t(8;21)(q22;22) |
| molecular alterations (from DepMap) | P15INK4B, P16INK4A, TP53 | NRAS | TP53, KRAS | TP53, NRAS | P16INK4A, TP53, KRAS, RB1 | RAD21, TP53, KIT | TP53, KIT |

**Supplementary Table 9:** Information about patient derived xenograft cells

| Figure ID | ML-DS PDX | AMKL PDX |
|---|---|---|
| sex | m | m |
| age at diagnosis | 1y 4m | 0y 11m |
| WBC (x$10^9$/L) | 32.5 | 23 |
| hemoglobin (g/dl) | 7.9 | 11.8 |
| BM blasts (%) | 71 | 46 |
| CNS | No | No |
| SCT | Yes | Yes |
| molecular mutations | GATA1 | KMT2A |
| Cytogenetics (karyotype) | 47,XY,t(3;13)(q?26;q?13~14),del(13)(q?14q22),+21c[cp14]/47,sl,del?(15)(q?)[cp2]/46,XY[1] | 46,XY[15].nuc ish,3q26(EVI1x2)[100/100],8q22(RUNX1T1x2),21q22(RUNX1x2)[98/100],11q23(MLLx2)[99/100],16q22(CBFBx2)[100/100],17q21.1(RARAx2)[100/100] |
| response | NR, CCR after SCT | CCR |
| relapse | No | Yes |
| Used in CFU assays | Yes | Yes |
| Used in transplants | Yes | Not yet |

**Supplementary Table 10:** sgRNA sequences

| **Name** | **Primer sequence** |
|---|---|
| sg1_CRISPRi (sgHME-1) | GCAGTACTTTGGTAAGTGTG |
| sg2_CRISPRi | AGGTTTGCTAATGGGATGAG |
| sg3_CRISPRi (sgHME-2) | GTAAGTGTGAGGCTGTCTCA |
| sg4_CRISPRi | CTGAAACCAAAGCAGTACTT |
| lncHME_sg1_CasRX | CTTTAGAGAACGTAGCCTCACCAGTCATCA |
| lncHME_sg2_CasRX | TAATACATATTATAACACCACACTGTGAAA |
| lncHME_sg3_CasRX | GATTGCTCATAAGACCTACAGTCAGAAAGG |
| lncHME_sg4_CasRX | TGGTATTACGATGACATGCTCCAATTTCTT |
| sgRNA_247_tiling | AGGGGTGGTGGGAACACAGC |
| sgRNA_725_tiling | GGGGTGGTGGGAACACAGCA |
| sgRNA_738_tiling | GGTGGTGGGAACACAGCAGG |
| sgRNA_736_tiling | GGTGGGAACACAGCAGGAGA |
| TAL1_sg3 (sgTAL1) | GTATGAGATGGAGATTACTGA |
| GATA1_sg1 (sgGATA1) | GCCATTGCTCAACTGTATGGA |

**Supplementary Table 11:** qPCR primer

| Name | Primer sequence |
|---|---|
| FW_lncHME_qPCR | TGCAGAAGAGCAGCTGAAAC |
| RV_lncHME_qPCR | TCCCAAGGATGCTGGTTGTAG |
| FW_AHI1_E24-25_qPCR | AGATACAGCACCAACGGTAGTG |
| RV_AHI1_E24-25_qPCR | TCCCTTTCCTATGCTGCCATAC |
| FW_AHI1_E27-28_qPCR | CAGGACTTCAGACTAGGCTCAG |
| RV_AHI1_E27-28_qPCR | TTCTGCCTGCTTGCTTGTTC |
| FW_MYB_qPCR | GGGAACAGATGGGCAGAAATCG |
| RV_MYB_qPCR | GCTGGCTTTTGAAGACTCCTGC |
| FW_HBS1L_qPCR | TGCATGATGAACCTGTCGAC |
| RV_HBS1L_qPCR | TGGGGCCACAAAATATGCAG |
| FW_TAL1_qPCR | CCACCAACAATCGAGTGAAGAGG |
| RV_TAL1_qPCR | GTTCACATTCTGCTGCCGCCAT |
| FW_GATA1_qPCR | CATCCGGCCCAAGAAGCGCC |
| RV_GATA1_qPCR | CGCATGGTCAGTGGCCGGTT |
